# Integrative single-cell characterization of hypothalamus sex-differential and obesity-associated genes and regulatory elements

**DOI:** 10.1101/2022.11.06.515311

**Authors:** Hai P. Nguyen, Candace S.Y Chan, Dianne Laboy Cintron, Rory Sheng, Lana Harshman, Mai Nobuhara, Aki Ushiki, Cassidy Biellak, Kelly An, Gracie M. Gordon, Francois Mifsud, Abbey Blake, Eric J. Huang, Martin Hemberg, Christian Vaisse, Nadav Ahituv

## Abstract

Over 500 noncoding genomic loci are associated with obesity. The majority of these loci reside near genes that are expressed in the hypothalamus in specific neuronal subpopulations that regulate food intake, hindering the ability to identify and functionally characterize them. Here, we carried out integrative single-cell analysis (RNA/ATAC-seq) on both mouse and human male and female hypothalamus to characterize genes and regulatory elements in specific cell subpopulations. Utilizing both transcriptome and regulome data, we identify over 30 different neuronal and non-neuronal cell subpopulations and a shared core of transcription factors that regulate cell cluster-specific genes between mice and humans. We characterize several sex-specific differentially expressed genes and the regulatory elements that control them in specific cell subpopulations. Overlapping cell-specific scATAC peaks with obesity-associated GWAS variants, identifies potential obesity-associated regulatory elements. Using reporter assays and CRISPR editing, we show that many of these sequences, including the top obesity-associated loci (*FTO* and *MC4R*), are functional enhancers whose activity is altered due to the obesity-associated variant and regulate known obesity genes. Combined, our work provides a catalog of genes and regulatory elements in hypothalamus cell subpopulations and uses obesity to showcase how integrative single-cell sequencing can identify functional variants associated with hypothalamus-related phenotypes.

## Introduction

Obesity dramatically increases the risk of hypertension, dyslipidemia, diabetes, COVID-19, and cardiovascular diseases and is a major public health concern^1,2^. Environmental and genetic factors are both involved in the onset and progression of weight gain^3^, with genetic factors being a major component^4,5^. Both rare, monogenic obesity causing variants, and common variants, detected through large obesity genome wide association studies (GWAS), have been found to predispose to obesity. While rare gene mutations causing obesity have outlined a neuro-physiological pathway involved in body weight regulation, the majority of which are part of the hypothalamic leptin-melanocortin pathway^6^, how common variants influence body weight remains poorly understood. GWAS have identified over 500 common loci associated with obesity^6-9^. The majority of obesity GWAS loci represent clusters of common variants in noncoding regions that likely alter gene expression and have been found to reside primarily near genes involved in central nervous system (CNS)-related processes^9-11^. In addition, a number of loci appear to directly influence neurons of the hypothalamic leptin-melanocortin system^10,12^. Combined, these studies single out neuronal subpopulations in the hypothalamus that are involved in food intake regulation as the major cause of this condition.

The hypothalamus is one of the most complex regions in the brain, composed of diverse neurons and glial cells that are spatially distributed in different regions or nuclei^13-15^. Extensive studies of the hypothalamus have demonstrated that individual nuclei have distinctive functions in regulating different homeostatic processes, such as thermoregulation, mating motivation, sweat, blood pressure, reward, sleep and hunger^16-22^. Most of these processes are controlled by neuronal subtypes in each nucleus. In addition, sex-dimorphism in the hypothalamus has been characterized^23-25^, with the medial preoptic nucleus being a major sex-dimorphic region, releasing different hormones depending on sex^26-28^. However, the understanding of sex-differential gene expression and transcriptional regulation in the hypothalamus remains largely uncharacterized.

With advances in single-cell sequencing technologies, several studies have set out to characterize diverse cell populations in the whole hypothalamus or specific nuclei using single-cell RNA sequencing (scRNA-seq)^29-41^. These studies characterized various hypothalamus neuronal populations, such as GABAergic and glutamatergic neurons, non-neuronal cell types, including microglia, oligodendrocytes, and astrocytes. While these studies were able to identify various cell populations based on their transcriptional profiles, they lack the capacity to identify the regulatory elements driving these cell type-specific subpopulations, where many variants of the obesity GWAS or other hypothalamus-associated phenotypes could reside.

Integrative single-cell sequencing utilizes RNA-seq to identify the transcriptome and ATAC-seq to characterize the regulome in the same cell. Here, we utilized this technique to generate a combined scRNA- and sc-ATAC seq atlas of the adult female and male mouse and human hypothalamus. This atlas, composed of more than 50,000 cells, identified numerous non-neuronal and neuronal cells that play key roles in controlling various hypothalamus-associated physiological processes, including blood pressure, mating, nursing, and feeding. We also identified cell-type specific regulatory elements and linked them to their target genes in several cell clusters. Moreover, we uncover cell cluster-specific transcription factor (TF) activity and their regulated genes and identify cell cluster-specific core TFs shared between mice and humans. In addition, we annotated sex-associated gene expression and regulatory element differences, including hormones/neuropeptides in different neuronal subtypes that are regulated by TF expressed in a sex-dimorphic manner. Overlapping our cell-type specific regulatory elements with obesity GWAS variants, we identified eighteen potential enhancers for several obesity associated loci, including *FTO* and *MC4R*, the top two associated obesity variants^42,43^. Enhancer assays in mouse hypothalamus cells, identify ten of these to be functional enhancers and five to alter enhancer activity due to the obesity-associated variant. Genetic disruption or CRISPR inactivation (CRISPRi) mediated downregulation of these enhancers in cultured human neurons or astrocytes leads to a reduction of gene expression of their target genes. Taken together, this work provides a catalog of genes and regulatory elements in different cellular subpopulations of the hypothalamus in both males and females in mice and humans which can be utilized to identify causative sequences that lead to hypothalamus-associated phenotypes, such as obesity.

## Results

### Combined sc-RNA and ATAC profiling of the mouse hypothalamus

To characterize both the genes and regulatory elements of various hypothalamic cell types, we utilized the 10X Chromium multiome platform. We initially focused on mouse, utilizing three male and three female hypothalami dissected from 6 weeks old C57BL/6J mice (**Fig.1a**). We isolated nuclei via FACS and subjected them to 10X Chromium Multiome following established protocols (see **Methods**). Overall, we sequenced 72,504 cells. After quality control filtering (see **Methods**), we retained 13,455 from female and 13,329 from male mice, amounting to 26,784 mouse hypothalamus cells in our dataset (**Extended Data Fig.1a**). We next set out to characterize the various hypothalamus cell populations using our scRNA-seq data. Utilizing differential expressed genes and cell-type specific markers curated from literature and previous studies in hypothalamus, we first determined the various cell-type clusters^29-41^. We identified 37 distinctive cell populations, including 34 previously-described cell-types and four specialized neurons that were not described in previous studies. These include 33 different types of neurons and oligodendrocytes, astrocytes, and microglia cell clusters (**Fig.1b**).

**Fig 1.**
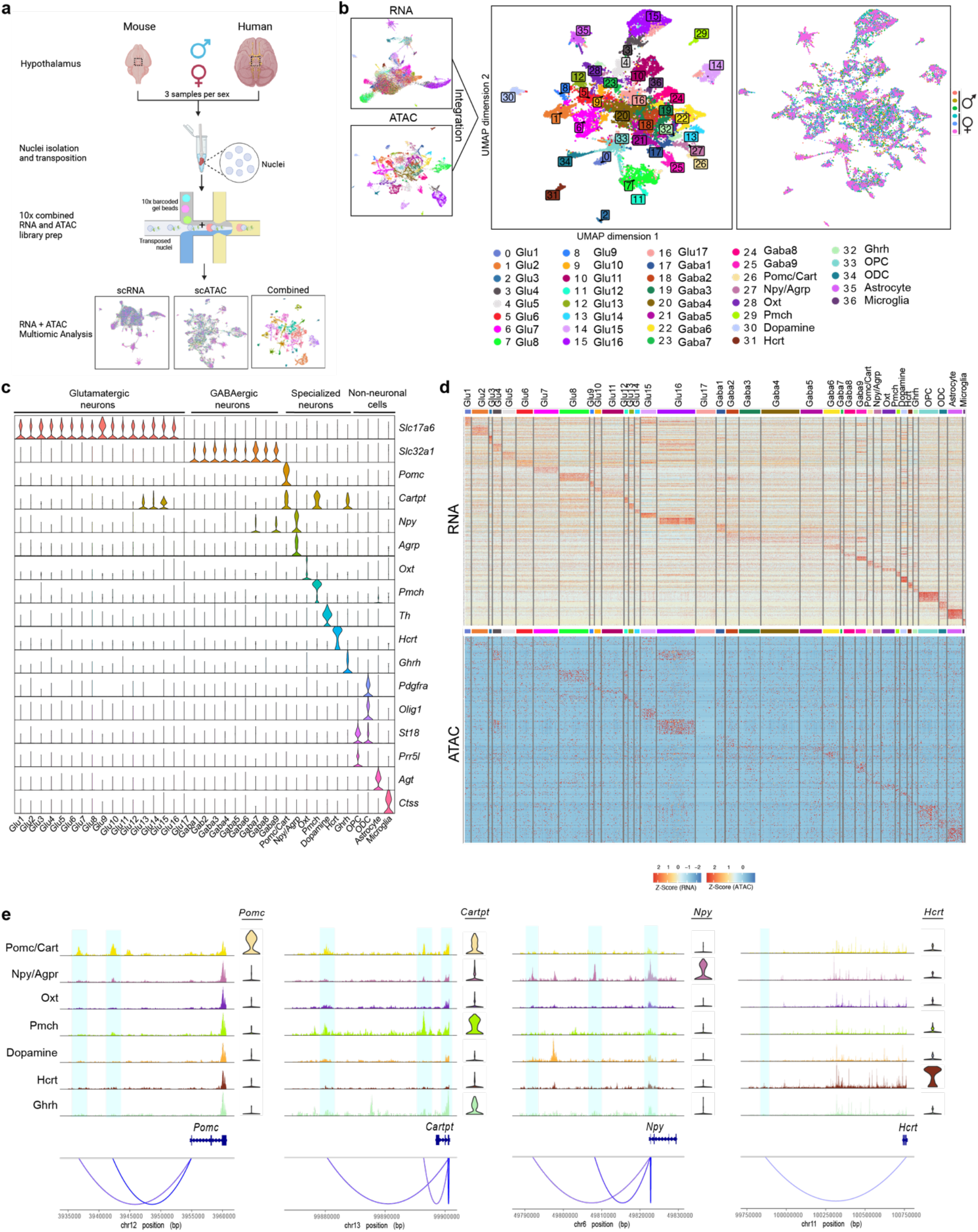
Integrative single-cell RNA and ATAC atlas of adult mouse hypothalamus. **a**, Schematic of preparation of mouse and human hypothalamus of three male and three female samples and 10X multiomic library prep. **b**, Uniform Manifold Approximation and Projection (UMAP) of single-cell RNA, ATAC, and integrative data (center). Clusters were identified based on scRNA-seq data. Right panel shows the UMAP by subject. **c**, Violin plot of gene markers used to identify all clusters. **d**, Heatmap of the top five differentially expressed genes and the top ten ATAC peaks across all cell populations. **e**, scATAC peaks to gene linkage plots for *Pomc, Cartpt, Npy*, and *Hcrt* with genome tracks of specialized neuron populations, including Pomc/Cart, Npy/Agrp, Oxt, Pmch, Dopamine, Hcrt, and Ghrh cell clusters. Gene expression violin plots for each cell type are shown on the right. The linked scATAC peaks to gene promoters are highlighted in light blue.

For neuronal cells, we found 33 different neuronal cell populations, including glutamatergic, GABAergic, and specialized neuron clusters. We found 17 glutamatergic neuron populations that showed high expression levels of the glutamatergic marker, *Slc17a6*, which encodes vesicular glutamate transporter 2 (*Vglut2*). We also identified nine clusters of GABAergic neurons, expressing GABAergic marker *Slc32a*, which encodes the vesicular GABA transporter (*Vgat*) (**Fig.1c**). The GABAergic neurons also expressed the synthetic enzyme for GABA, *Gad1* (**Extended Data Fig.1b**). Furthermore, we found other specialized neurons secreting various neuronal peptides and hormones that regulate whole-body endocrine and physiological homeostasis. Among these cells, we identified Pomc/Cart neurons that distinctly express both proopiomelanocortin (*Pomc*) and cocaine and amphetamine regulated transcript (*Cartpt*) and Npy/Argp neurons, highly expressing neuropeptide Y (*Npy*) and agouti-related peptide (*Agrp*) (**Fig.1c**). Both cell types are known to control satiety by regulating melanocortin 4 receptor (*Mc4r*) neurons activity^44^ (**Extended Data Fig.1b**). These neurons were identified in various nuclei, including paraventricular nucleus, accurate nucleus, and intermediolateral nucleus^45,46^. Similar to previous reports^47,48^, we found that Pomc/Cart and Npy/Agrp neurons express leptin receptor (*Lepr*) (**Extended Data Fig.1b**). In addition, we identified oxytocin (*Oxt*)- and pro-melanin concentrating hormone (*Pmch*)-secreting neurons (**Fig.1c**). While *Oxt* is known for its role in female reproduction, breastfeeding, and childbirth^49^, *Pmch* controls skin pigmentation and also has important roles in regulating motivated behaviors, such as feeding, drinking, and mating^50^. We also identified dopamine-producing neurons, which show high expression levels of dopamine synthetic enzymes, such as tyrosine hydroxylase (*Th*) and dopa decarboxylase (*Ddc*) (**Fig.1c, Extended Data Fig.1b**). We annotated hypocretin neuropeptide (*Hcrt*) or orexin -expressing neurons, which are reported to regulate arousal, wakefulness, and appetite^51,52^. Additionally, we found growth hormone-releasing hormone (*Ghrh*)-specific neurons (**Fig.1c**), which regulate growth hormone production in the anterior pituitary gland^53,54^. Analysis of other neuronal peptides, found oxytocin neurons expressed *Avp* (vasopressin), regulating vascular tone, blood pressure and urine concentration by the kidney^55^ (**Extended Data Fig.1c**). We found that *Ghrh* neurons highly expressed *Gal* (Galanin-like peptide), which is known to regulate feeding and reproduction^56^ (**Extended Data Fig.1c**). In addition, glutamatergic neurons widely expressed *Adcyap1*, encoding for pituitary adenylate cyclase-activating polypeptide binding to its receptor in intestinal cells and this gene is associated with post-traumatic stress disorder (PTSD)^57^ (**Extended Data Fig.1c**).

We also identified several non-neuronal cell populations. These include oligodendrocytes (ODC), oligodendrocyte precursor cells (OPC), which show high expression of suppression of tumorigenicity 18 (*St18*) and proline-rich 5 like (*Prr5l*) genes and mature oligodendrocytes, expressing platelet-derived growth factor receptor A (*Pdgfra*) and oligodendrocyte transcription factor 1 (*Olig1*) (**Fig.1c**). In addition, we found astrocytes that show high levels of astrocyte gene markers, such as angiotensinogen (*Agt*) expression and microglia that express the immune marker cathepsin S (*Ctss*) (**Fig.1c**).

We next examined our scATAC-seq data, obtained from cells that were jointly profiled with scRNA-seq. Dimensionality reduction was performed using latent semantic indexing and batch correction using Harmony^58^ (see **Methods**). Peak calling on each cell-type from joint clustering recovered 359,963 chromatin accessible peaks from cells that passed quality control. Amongst them, 157,204 (43%) peaks were found in only one cluster, indicating robustness of clustering and annotation. Comparing differential gene expression and accessible peak profiles, found that all our cell populations had unique gene and chromatin signatures in a cell-specific manner (**Fig.1d**). We found 37,556 marker peaks that were enriched in specific cell-types, with many of them located near cell type-specific gene markers (5,400). For example, we found increased accessibility near *Slc176* in glutamatergic neuron cell types, *Slc32a6* in GABAergic neuronal populations, *Ddc* in dopamine neurons, and *Hcrt* in Hcrt neurons (**Extended Data Fig.1d**).

Combined scATAC and scRNA sequencing allows to identify potential *cis*-regulatory elements by annotating peaks where chromatin accessibility correlates with gene expression (see **Methods**). We found 245 peaks that were linked to the expression of 65 marker genes. Among the specialized neurons, *Pomc* expression was linked to two scATAC-seq peaks that were highly specific to Pomc/Cart neurons cells (**Fig.1e**). These elements overlapped with two enhancers reported in^59^. We also identified cell type-specific scATAC-seq peaks that were linked to *Cartpt* expression in neuronal populations, such as Pomc/Cart and *Pmch* neurons that highly express *Cartp* (**Fig.1e**). In addition, we annotated three *Npy* linked scATAC-seq peaks that were highly specific to *Npy*/*Argp* neurons. *Pomc* and *Cartpt* are key regulators of feeding, metabolic homeostasis and obesity^60,61^ and their enhancer identification could be used to modulate their expression. We also found a *Hcrt* neuron specific scATAC-seq peak that was linked to *Hcrt* expression (**Fig.1e**) and a specific peak that was linked to *Th* expression in dopamine neurons (**Extended Data Fig.1e**). Moreover, there were multiple peaks that were linked to *Slc17a6* expression in GABAergic neurons (**Extended Data Fig.1e**). Combined, these results show the ability of integrative single-cell analyses to identify specific hypothalamus cell subpopulations and link potential gene regulatory elements with their putative target genes.

### Combined sc-RNA and ATAC profiling of the human hypothalamus

We next conducted a similar integrative single-cell experiment on three males and three female adult human hypothalami. We collected post-mortem hypothalami from patients ages 43-82 and a post-mortem interval (PMI) between 4-12 hours (**Supplementary Table 1, Extended Data Fig.2a**) and carried out scRNA/ATAC-seq, as described for mice. Due to the longer PMI of these samples, compared to mice, we initially obtained a lower number of cell populations. We thus carried out an additional scRNA/ATAC-seq experiment on the same samples subjecting them to NeuN (neuronal marker) FACS sorting to increase for neuronal populations. We combined both experiments and sequenced a total of 113,854 cells, which after filtering for high quality cells retained 34,257 cells (see **Methods**; **Extended Data Fig.2b**).

To improve the quality of our analyses, we removed estimated ambient RNA from our data using SoupX^62^. Joint analysis of scATAC and scRNA identified five broad clusters, which we annotated as mature oligodendrocytes, oligodendrocyte progenitor cells, astrocytes, microglia, and neurons (**Fig.2a**). Using differentially expressed genes, we found that neuronal clusters showed high expression for excitatory neuronal markers, including *GAD2*, which encodes for glutamate decarboxylase 2 and *NRG1*, encoding neuregulin 1 (**Fig.2b**). They also expressed other neuron markers, including stathmin 2 (*STMN2*) and synaptotagmin (*STY1*). Astrocytes expressed astrocyte gene markers, such as glial fibrillary acidic protein (*GFAP*) and *AGT* (**Fig.2b**). Similar to mice, OPC expressed oligodendrocyte markers, such as *PDGFRa* and chondroitin sulfate proteoglycan 4 (*CSPG4*). More mature oligodendrocytes expressed proteolipid protein 1 (*PLP1*) and myelin oligodendrocyte glycoprotein (*MOG*) (**Fig.2b**). We also found microglia cells with high expression of the pan-immune marker *PTPRC* (CD45), complement C1q B chain (*C1QB*), integrin alpha M (*ITGAM*), and integrin subunit alpha X (*ITGAX*) (**Fig.2b**). We found distinctive gene expression and scATAC profiles that are highly specific to each cell type (**Extended Data Fig.2c**).

**Fig 2.**
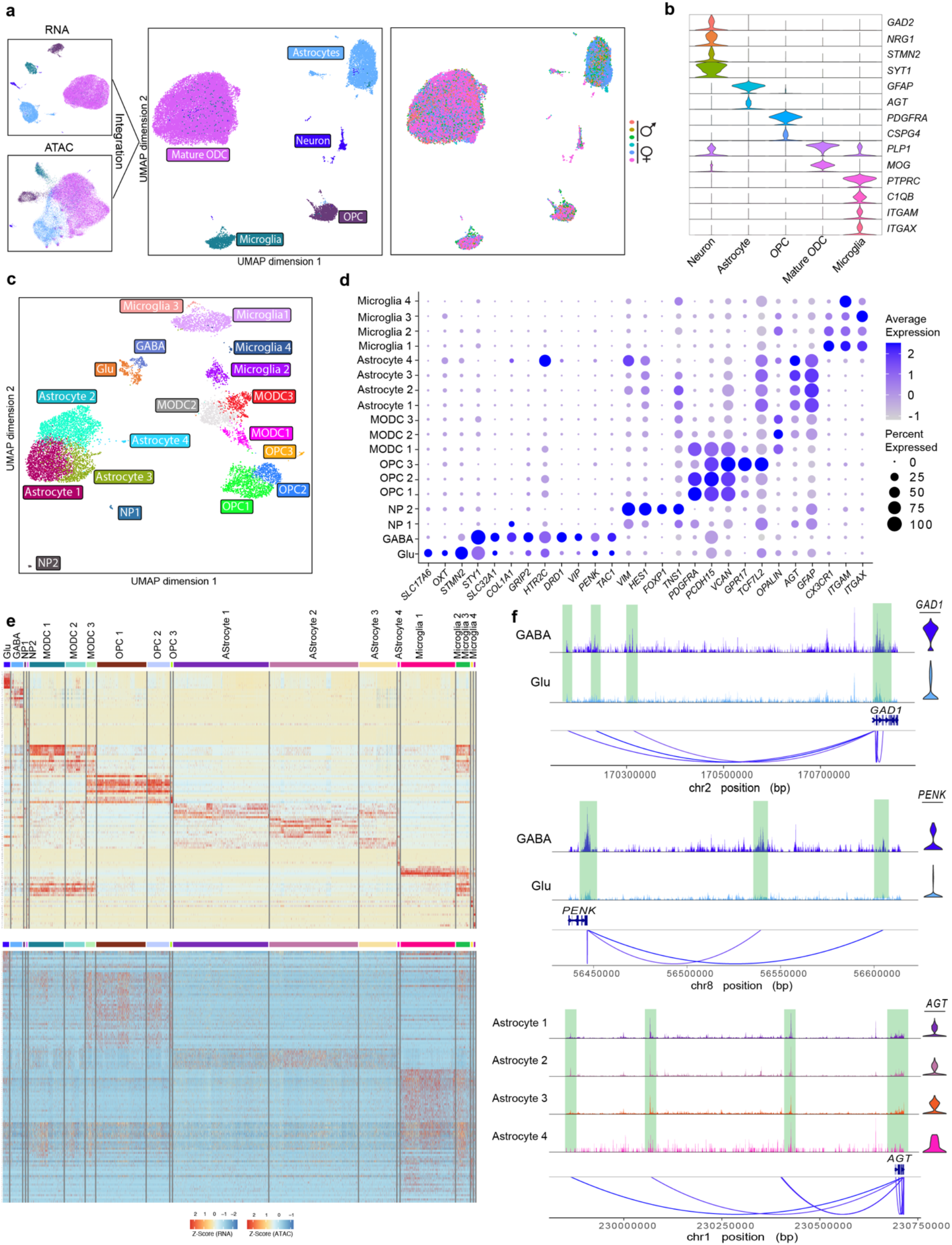
Integrative single-cell RNA and ATAC atlas of adult human hypothalamus. **a**, Uniform Manifold Approximation and Projection (UMAP) of single-cell RNA, ATAC, and integrative data of human hypothalamus samples. Right panel shows the UMAP by subject. **b**, Violin plot of gene markers used to identify cell clusters. **c**, UMAP of integrative data after removal of mature oligodendrocytes. Clusters were identified based on scRNA-seq data. **d**, Dot plots of gene markers used to identify cell clusters. **e**, Heatmap of the top five differentially expressed genes and the top ten ATAC peaks across all cell populations. **f**, scATAC peaks to gene linkage plots for *GAD1, PENK*, and *AGT* in GABAergic and glutamatergic neurons and astrocytes with the genome tracks. Gene expression violin plots are shown on the right. The linked scATAC peaks to gene promoters are highlighted in light green.

Despite the neuron top-off experiment, we found that the majority of cells (69.7%) isolated from human samples consisted of mature oligodendrocytes and had a much smaller neuronal population (1.7%). This low neuronal population recovery is consistent with other recent multiomic single-cell assays on post-mortem human midbrain or frontal lobe^63,64^. To increase the resolution of neuronal cells and other smaller populations, we subclustered these cells without the mature ODC, retaining 10,356 cells. Removal of the ODC cluster and re-clustering led to the identification of GABAergic and glutamatergic neuronal clusters, comprising 145 cells and 271 cells respectively (4.01%). In addition, we found two neural progenitor clusters, three astrocyte clusters, four microglia clusters, three myelinating oligodendrocytes (MODC) and three OPC clusters originating from the OPC population (**Fig.2c**). After subsetting our dataset, we found that 38.2% of peaks were specific to one cluster.

Similar to the mouse data, glutamatergic neurons had high expression levels of *SLC17A6* while GABAergic neurons showed high *SLC32A1* expression (**Fig.2d**). In glutamatergic neurons, we found oxytocin-expressing neurons with high expression of *OXT* and *AVP* (**Fig.2d, Extended Data Fig.2d**). We also found *POMC*- and *CARTPT*-expressing neurons, as well as *NPY*-expressing cells (**Extended Data Fig.2d**) and *PMCH, HCRT*, and *GHRH* neurons (**Extended Data Fig.2d**). GABAergic neurons showed high expression levels of *COL1A1, GRIP2*, and hydroxytryptamine receptor 2c (*HTR2C*), and dopamine receptor D1 (*DRD1*) (**Fig.2d**). In the GABAergic neuronal population, we also found neurons expressing neuropeptides, such as vasoactive intestinal peptide (*VIP*), proenkephalin (*PENK*) which plays a role in pain perception and response to stress^65^, tachykinin (*TAC1*) which is a ghrelin target and is involved in energy balance and food intake^66^, and thyrotropin releasing hormone (*TRH*) which stimulates the release of thyroid stimulating hormone (*TSH*) and prolactin from the anterior pituitary^67^ (**Extended Data Fig.2d**). Interestingly, we identified two neural progenitor (NP) clusters that showed high expression of neural progenitor gene marker, Vimentin (*VIM*) and stem cell/progenitor marker, transcription factor *HES1* (**Fig.2d**).

Using markers from previous single-cell studies on oligodendrocytes^68^, we identified three OPC and three myelinating ODC (MODC) populations. We found that OPC clusters had high expression levels of *PDGFRA*, protocadherin related 15 (*PCDH15*) and versican (*VCAN*) (**Fig.2d**). As OPC differentiate to myelin oligodendrocytes, they expressed G protein-coupled receptor 17 (*GPR17*), a Gi-coupled GPCR, that acts as an intrinsic timer of oligodendrocyte differentiation and myelination^69^. The MODC population showed high expression of transcription factor 7 like 2 (*TCF7L2*) and oligodendrocytic myelin paranodal and inner loop protein (*OPALIN*) (**Fig.2d**). We also identified four astrocyte clusters with high expression of *AGT* and *GFAP*. Examination of gene expression profiles of these astrocyte clusters found that each had a unique function. Genes that were highly expressed in MODC1 were involved in glutathione transport, plasma membrane repair and mitochondria distribution. In contrast, the MODC2 cluster showed high expression for genes involved in amino acid transport. Genes expressed in MODC3 were involved in glutamine transport and detoxification of copper ions, cholesterol and inorganic compounds. In addition, MODC4 expressed genes were involved in response to salt stress and calcium transport (**Extended Data Fig.2e**). Among the non-neuronal cell types, we also identified four microglia populations with high *ITGAM* and *ITGAX* expression (**Fig.2d**). In addition, we found that CX3CR1 and CD86, gene markers for activate/proinflammatory microglia, were expressed most highly in the microglia 1 cluster and at lower levels in microglia 2 and microglia 3 clusters. Microglia cluster 4 showed high expression for CD163, a gene marker for quiescent/anti-inflammatory microglia^70^ (**Fig.2d, Extended Data Fig.2f**).

We observed that all our cell populations had unique gene and chromatin profiles in a cell-specific manner (**Fig.2e**). We next examined our scATAC-seq data and found 65,560 peaks (38.2%) peaks found in only one cluster. 1,271 were differentially accessible, of which 30.9% (394) of differentially accessible peaks were located near cell type-specific gene markers. For example, we found higher number of reads in ATAC peaks near *GAD2* and *OXT* in GABAergic neurons, *GFAP* in the astrocyte populations and *CDPG4* in the MODC clusters (**Extended Data Fig.2g**). We next analyzed 46 cell marker genes to see whether chromatin accessibility in their locus correlated with gene expression. We examined chromatin accessibility up to one million base pairs away from the marker gene, which is roughly the median topologically associated domain (TAD) length^71^. Amongst the 46 marker genes analyzed, we identified 490 regions where accessibility correlated with gene expression. These regions were on average 289,499 (median 189,142) base pairs away from the transcription start site (TSS) of their postulated target gene. For example, we found that the expression of *GAD1*, a GABAergic neuronal marker, was linked to four scATAC-seq peaks in GABAergic neurons (**Fig.2f**). We also found three scATAC-seq peaks linked to the expression of proenkephalin (PENK). In addition, in the four astrocyte clusters, we found the expression of angiotensinogen (*AGT*), which regulates blood pressure, to be linked to seven scATAC-seq peaks (**Fig.2f**). Similar to our mouse data, using the integrative single-cell analyses allowed us to identify specific cell subpopulations in the human hypothalamus and potential gene regulatory elements with their putative target genes.

### Motif enrichment analyses identify cell-type specific regulatory pathways

We next set out to identify the major transcription factors (TFs) that regulate the various hypothalamus cell types, initially starting with mice. We identified motifs enriched in accessible regions for each cell-type using chromVAR^72^, finding distinct profiles of TF binding sites (TFBS) for each cell cluster. For example, for the glutamatergic neuron populations, we found significant TFBS enrichment of hypothalamic transcription factor nescient helix-loop-helix 2 (NHLH2), a known regulator of metabolic activity and puberty in the paraventricular nucleus (PVN)^73,74^(**Fig.3a**). We also found enrichment for the forkhead box P1 (FOXP1) motif and the NeuroD family of basic helix-loop-helix 1 (NEUROD1) motif, both known to regulate glutamatergic neurogenesis^75,76^(**Fig.3a**). In addition, we found the TFBS for cutlike homeobox 1 (CUX1), which controls neuronal proliferation, dendrite branching and synapse formation to be enriched in GABAergic neurons^77^(**Fig.3a**). Dopamine neurons showed enrichment for FOXA1/FOXA2 motifs, known to be associated with dopamine biosynthesis^78^. For non-neuronal populations, we identified binding motif enrichment of known cluster-specific TFs, such as *SOX9* in OPCs^79^, *BHLH21*, encoded by *Olig1* in oligodendrocyte and Spi-B transcription factor (Spi-1/PU.1 related; *SPIB*) known to regulate microglia/macrophages^80^(**Fig.3a**).

**Fig 3.**
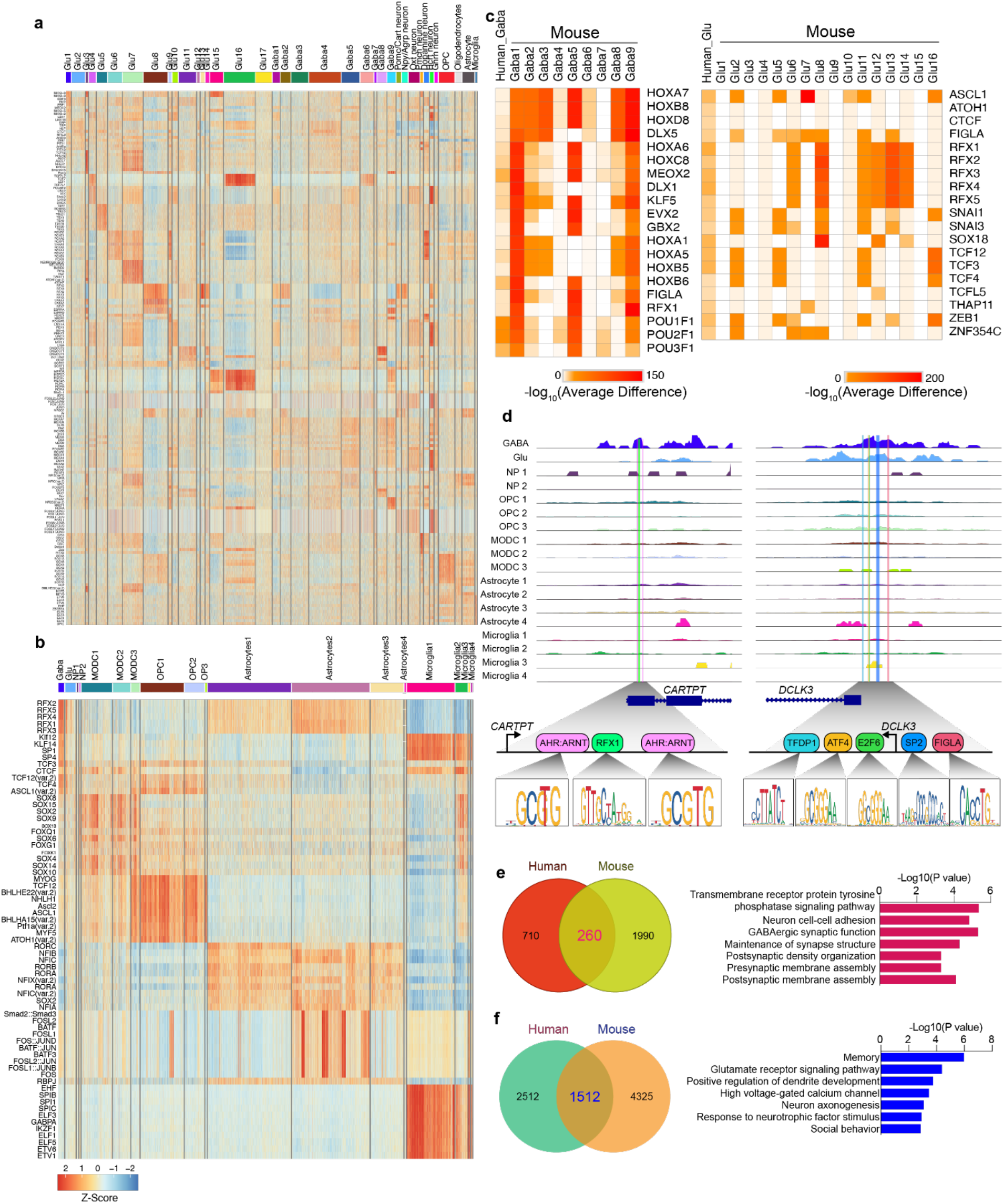
Motif enrichment analyses identify cell-type specific regulatory pathways. **a-b**, Heatmap of the top five differentially enriched transcription factors binding sites (TFBS) across cell populations labeled with different colors in the mouse (**a**) or human (**b**) hypothalamus. **c**, Heatmap of the top TFBS enriched in GABAergic and glutamatergic neuronal populations in mouse and human hypothalamus. **d**, Genome tracks of *CARTPT* and *DLCK3* showing logo plots for various TFBS in their promoters. **e-f**, Venn diagrams showing the overlap of TFBSs enriched in GABAergic (**e**) or glutamatergic (**f**) neurons of mouse and human hypothalamus along with gene ontology analysis of enriched TFBS in these neurons.

In humans, similar to mice, we identified unique TFBS profiles in a cell type-specific manner. For glutamatergic and GABAergic neurons, we observed an enrichment for the NHLH2 TFBS (**Fig.3b**). We found that both mouse and human neuronal populations have highly enriched motifs for regulatory factor binding to the X-box (RFX) family, including *RFX1* and *RFX3. SOX9* TFBS and its family members were enriched in OPC and MODC populations (**Fig.3b**). We also found a TFBS enrichment for transcription factor 3 (*TCF3*), which is known to maintain OPC^81^. In addition, we found *BHLH21* TFBSs were highly enriched in astrocytes along with nuclear factor IA (NIFA), which is known to induce astrocyte formation^82^ (**Fig.3b**). Similar to mouse data, microglia cells showed an enrichment for *SPIB* TFBSs (**Fig.3b**).

We next compared mouse and human TFBSs whose motifs were enriched in a cell-cluster specific manner. In both mouse and human GABAergic and glutamatergic neuronal populations, we observed a large number of shared TFBSs, such as *HOXA7, MEOX2, NHLH2, RFX1* and *RFX3* (**Extended Data Fig.3a,b**). Analysis of mouse and human GABAergic neurons for the TFBSs that have the most enriched motif scores, found many shared TFBS to be enriched in these neurons, such as distal-less homeobox 5 (*DLX5*), well-known to regulate GABAergic neurons and control behavior and metabolism^83^, even-skipped homeobox 1 (*EVX1*), which is required for GABAergic neuron development^84^, and *POU3F1* which regulates neural differentiation^85^(**Fig.3c**). In the glutamatergic neuronal populations, we found mouse and human cells to show TFBS enrichment for the RFX TF family, achaete-scute family BHLH transcription factor 1 (*ASCL1*), that is required for dopaminergic neurons differentiation from iPSC cells^86^ and *THAP11* or RONIN, known regulators of neural stem cell differentiation^87^(**Fig.3c**). In the glutamatergic neurons, we also found TFBS enrichment for TFs that have not been reported to have specific function in neuronal cells, such as folliculogenesis specific BHLH transcription factor (*FIGLA*), which has a known function in postnatal oocyte-specific gene expression and zinc finger E-box binding homeobox 1 (*ZEB1*), which is known to suppress the T-lymphocyte-specific IL2 gene^88^ (**Fig.3c**). In addition, we found that many other non-neuronal clusters in mouse and human hypothalamus shared many TFBSs that potentially regulate genes in these cell types (**Extended Data Fig.3c**). Combined, these results suggest that hypothalamus cell populations in mice and humans utilize a similar cell type-specific regulatory machinery.

We next tested whether the TFBS that were enriched in specific cell clusters are associated with TFs expression in the corresponding clusters. We compared TFBS enrichment to the corresponding TF expression, finding several TFs to be both differentially expressed and enriched in chromatin accessible regions (**Extended Data Fig.3d**). However, we did not find statistically significant correlation (r > 0.5, Pearson correlation) between TF expression and motif enrichment, possibly as TF concentration is not the sole determinants of DNA binding^89^. However, a small number of TFs which had enriched TFBS in a cluster-specific manner had high expression in the corresponding clusters. For example, *KLF5, FIGLA* and *RFX1* were found to be highly expressed and have enriched TFBS scores in GABAergic neurons (**Fig.3d**). We also found *THAP11* TFBS to be enriched and with high expression in glutamatergic neurons.

We also examined TFBSs corresponding to scATAC peaks in the promoters of marker genes of neuronal populations. Examination of the *CARTPT* promoter, a glutamatergic neuronal marker, found multiple TFBSs for hydrocarbon receptors (AHR:ARNT), which are known to regulate and determine neuronal fate^90^ and *RFX1* (**Fig.3d**), whose expression were high in glutamatergic neurons in mouse and rat^91,92^. In addition, we found multiple TFBSs for *ATF4, E2F6, FIGLA, SP2*, and *TFDP1* in the promoter of doublecortin like kinase 3 (*DCLK3*) (**Fig.3d**), which is highly expressed in GABAergic neurons (**Extended Data Fig.3e**) and is known to have a protective role in these cells^93^.

We next examined genes that might could be regulated by TFs with cell-type enriched TFBS in both mice and humans. For each TF, we searched for the nearest genes (within 2kb) of motif-containing cell-type accessible peaks (see **Methods**). We then compared the list of genes predicted to be regulated by TFs in neuron populations between humans and mice. In GABAergic neurons, we identified 970 regulated genes in humans and 2,250 genes in mice (**Fig.3e**). For GABAergic neurons, we found 260 genes that overlap between humans and mice, which following gene ontology (GO) analyses^94^ were found to be enriched for neuron-specific pathways, including neuron cell-cell adhesion, GABAergic synaptic function, and postsynaptic density organization (p-value < 0.05, hypergeometric distribution test) (**Fig.3e**). Interestingly, the human specific pathways in GABAergic neurons were found to be enriched for circulatory system development, cytoskeleton organization, and cell projection organization (**Extended Data Fig.3f**) while the mouse specific pathways in GABAergic neurons were enriched for cell-substrate adhesion, extracellular structure organization and cell-matrix adhesion (**Extended Data Fig.3g**). These results suggest that these species-specific pathways in GABAergic neurons are involved in general cell maintenance. In glutamatergic neurons, we identified 4,200 genes in humans and 5,837 genes in mice, with both species sharing 1,512 genes (**Fig.3f**). GO analysis of shared genes found them to be involved in memory, glutamate receptor signaling pathway and high voltage-gated calcium channels (**Fig.3f**). Human specific glutamatergic neuron-specific regulated genes were found to be involved in neuron projection, axon extension, and actin filament organization, while mice specific genes were involved in heparan sulfate proteoglycan biosynthesis, ERBB2 signaling pathway and response to forskolin (**Extended Data Fig.3h,i**). These data indicate that mouse and human hypothalamus cell populations share many common TFs that regulate similar cluster-specific gene networks and have networks unique to each species. In addition, our datasets provide the ability to identify TFs that are involved in cell-type specific hypothalamus gene-regulatory networks.

### Characterization of sex-differential gene and regulatory activity

The hypothalamus is a neuro-endocrine organ that releases various neuropeptides in a sex-specific manner. We thus annotated sex-differential gene expression across the cell populations in the human hypothalamus. We first identified genes that were differentially expressed between males and females in corresponding cell types. We detected sex-differential expressed genes across different cell types, but their numbers are small within some clusters (10 out of 18 clusters) having less than 10 sex-differentially expressed genes (**Fig.4a**). We next examined various neuropeptides in the neuronal populations. As a positive control^95,96^, we first examined the estrogen receptor (*ESR1*) and androgen receptor (*AR*), finding *ESR1* to be more highly expressed in glutamatergic neurons of females than males and *AR* to have higher expression in male GABAergic neurons (**Fig.4b**). When examining all neuropeptide genes, we found various genes to be differentially expressed in neuronal clusters between sexes. We found a number of neuropeptide genes to be expressed at higher levels in males than females, including *AVP, GHRH, GPR, NPY, HCRT*, and *TRH*. In contrast, we found *PENK* and *VIP* expression to be higher in females than males (**Fig.4b**). Analysis of these genes in mouse found them to be similarly sex-differentially expressed (**Extended Data Fig.4a**). Our findings are in line with previous studies. For example, *AVP* is more highly expressed in male GABAergic and glutamatergic neurons than in females and previous studies have shown that *AVP*, which regulates various social behaviors, including aggression and mating in males, has a higher expression in male hypothalamus than females^97,98^. We also found higher expression of *VIP* in female GABAergic neurons than males. In the hypothalamus, *VIP* is regulates the release of prolactin, stimulating milk production in the mammary gland^99,100^ and *VIP* expression is found to be higher in the female brain than males^100,101^. In addition, we found novel sex-differentially expressed neuropeptides, including *GHRH, GPR, HCRT*, and *TRH* (**Fig.4b**), whose physiological process has been indicated to be sex-dimorphic. However, their cell-type specific expression in neurons have not been reported. For example, growth hormone secretion has been demonstrated to be sex-dimorphic, and higher in males in both mice and humans^102^. Our data shows that *GHRH* (growth hormone releasing hormone) is more highly expressed specifically in glutamatergic neurons, leading to higher secretion of *GH* (**Fig.4b**).

**Fig 4.**
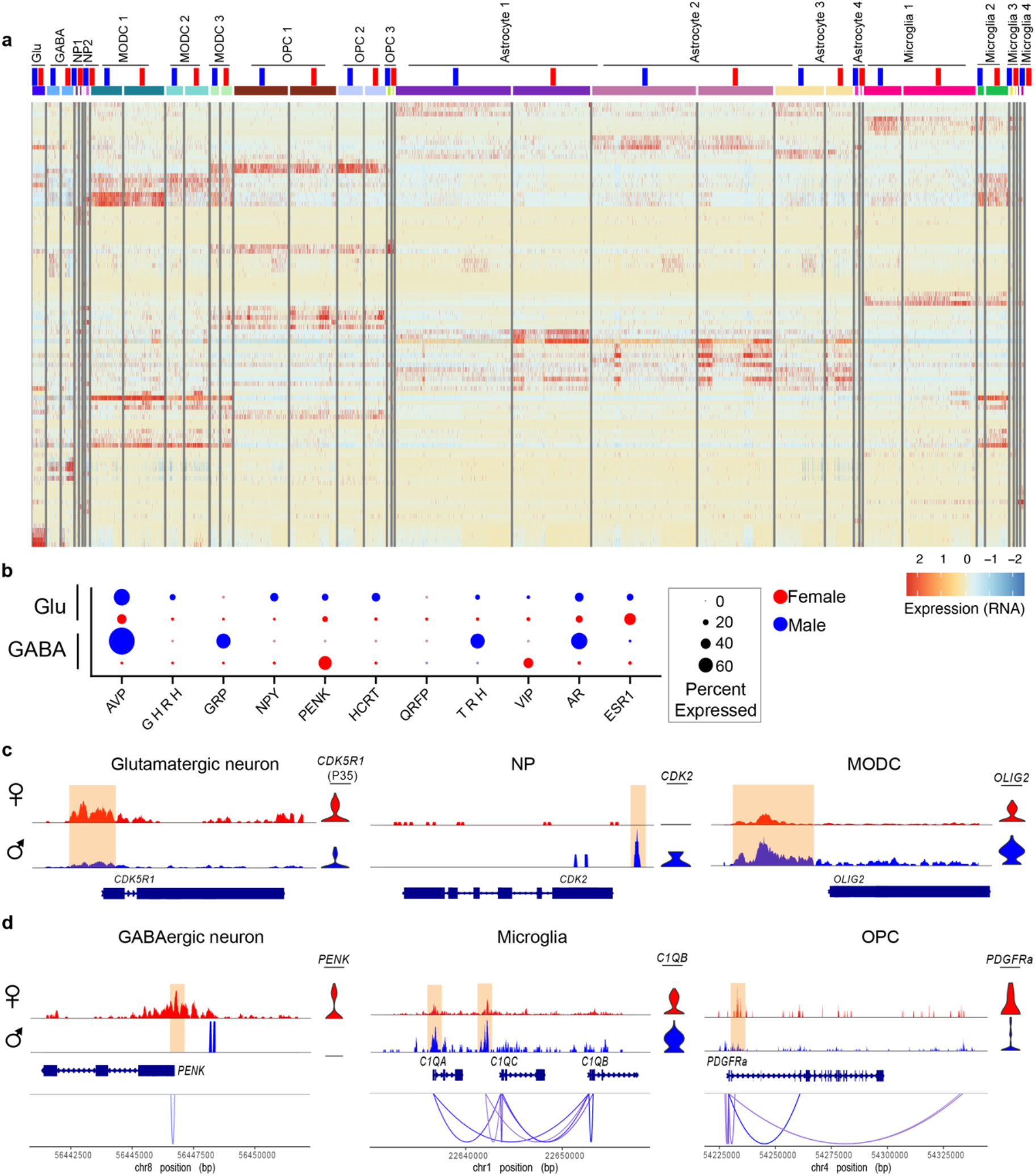
Human sex-differentially expressed genes and regulatory elements. **a**, Heatmap of the top five sex-differentially expressed genes for the corresponding cell populations in the human hypothalamus. Female cell populations are labeled in red and male cell populations in blue. **b**, Dot plot showing the expression of neuropeptides that were sex-differentially expressed in GABAergic and glutamatergic neurons. **c**, Genome tracks of sex-differentially expressed genes with sex-differential scATAC peaks at their promoters, including *CDK5R1* (P53) in glutamatergic neurons, *CDK2* in NP, and *OLIG2* in MODC clusters. **d**, Linkage plots showing sex-specific scATAC peaks linked to sex-differentially expressed genes, including *PENK* in GABAergic neurons, *C1QB* in microglia, and *PDGFRa* in OPC clusters.

To further dissect the molecular mechanism of sex-differentially expressed genes, we examined potential sex-differential accessible chromatin regions located in the promoters of the sex-differentially expressed genes. We found 54 sex-differentially expressed genes that had sex-differential scATAC peaks located near their promoters. For example, in glutamatergic neurons, we observed increased accessibility at the promoter region and increased gene expression for the cyclin dependent kinase 5 regulatory subunit 1 (*CDK5R1*)(**Fig.4c**), which encodes for P35, a neuron-specific activator of cyclin-dependent kinase 5 (*CDK5*) and is required for central nervous development^103^. In addition, cyclin-dependent kinase 2 (*CDK2*) had higher expression in female neural progenitor clusters, accompanied by a female-specific scATAC peak at its promoter (**Fig.4c**). We also found *OLIG2*, an oligodendrocyte gene marker and known activator of myelin-associated genes, to have higher expression in males and increased accessibility in its promoter in male ODC compared to females (**Fig.4c**). Oligodendrocytes are known to have proliferation and differentiation sex-differences, which leads to sex-dimorphic risk to neurodegenerative diseases^104^. Sex-differential expression of *OLIG2*, the key transcription factor of oligodendrocytes might be responsible for sex-dimorphic differentiation of oligodendrocytes.

We identified several sex-differential scATAC peaks linked to corresponding sex-differentially expressed genes. For example, in GABAergic neurons, a scATAC peak was linked to proenkephalin (*PENK*) expression in males, but not in females (**Fig.4d**). This gene plays a role in pain perception and response to stress and was shown to be more highly expressed in males versus female rats^105^, similar to what we observed in humans. In microglia populations, *C1QB*, which is involved in synaptic pruning, showed higher expression and stronger scATAC peaks in males versus females in line with previous analyses of the developing brain^106^. In addition, we found a scATAC peak that was linked to the expression of the platelet derived growth factor receptor alpha (*PDGFRa*), which plays an important role in cell growth and division, in the female OPC population (**Fig.4d**).

### Characterization of regulatory elements overlapping obesity-associated variants

We next set out to utilize our single-cell catalog of human hypothalamus cell-type specific genes and regulatory elements, to characterize obesity associated GWAS noncoding variants. We compiled a list of 508 index SNPs associated with obesity from the NHGRI-EBI GWAS catalog^107^. We performed linkage disequilibrium (LD) expansion on all the lead SNPs to include nearby variants with high probability of coinheritance (r^2^ >0.8). The majority of these SNPs (>97%) localize to noncoding regions of the genome. We then checked for overlap of these SNPs with scATAC-seq peaks (**Fig.5a**), finding 102 SNPs that overlap 35 marker peaks. These SNPs comprised some of the most highly associated loci with obesity, including the top two loci (*FTO* and *MC4R*)^108,109^. Among the 35 peaks, we had two scATAC peaks linked to *FAIM2*, two scATAC peaks linked to *FTO*, and two scATAC peaks linked to *MC4R*. We did not observe a specific cell-type that was more strongly associated with obesity, having overlapping scATAC peaks from various cell types including oligodendrocytes, oligodendrocyte precursor cells, astrocytes and neurons.

**Fig 5.**
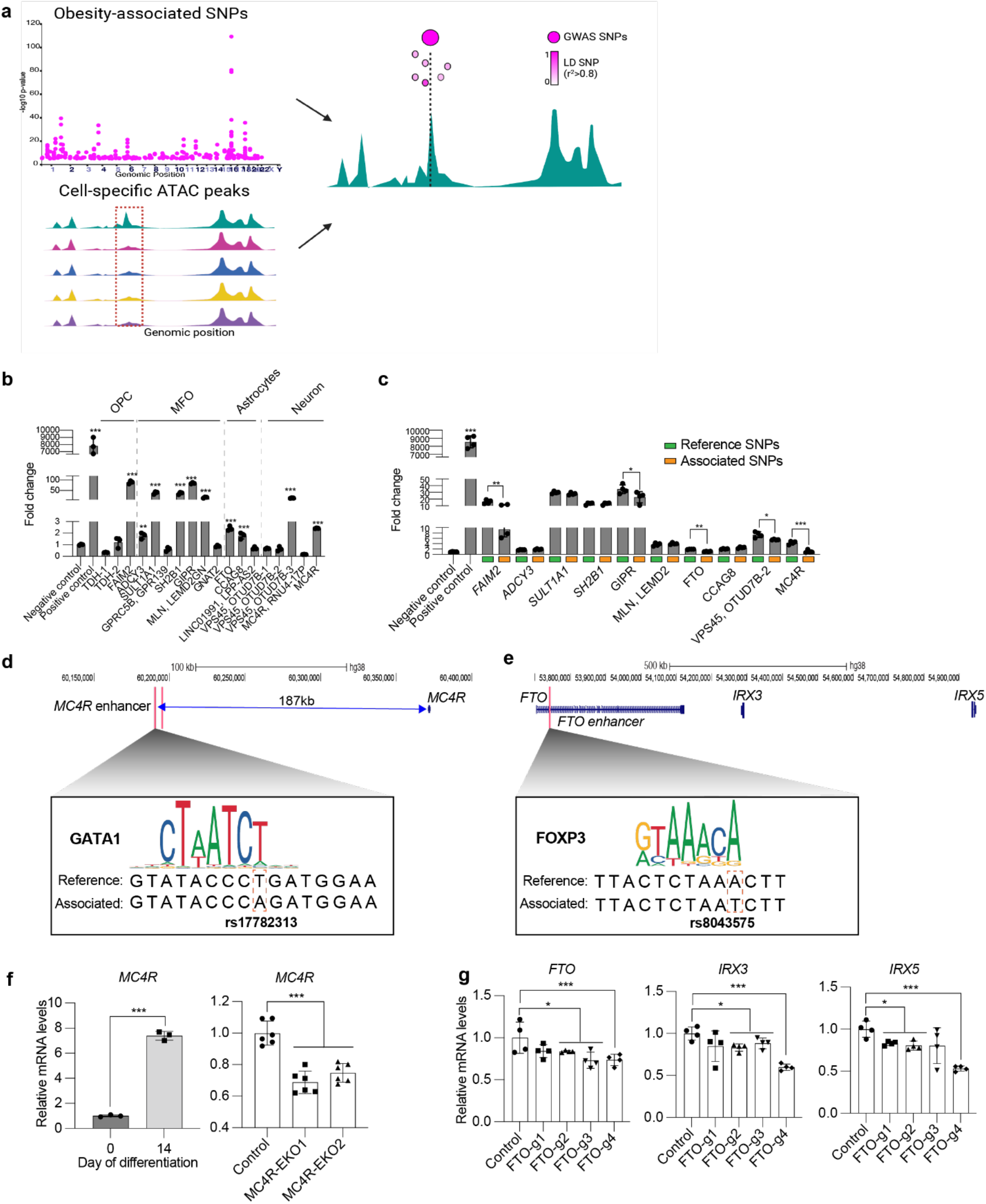
Integrative single cell sequencing data identifies cell-type specific gene regulatory elements overlapping obesity-associated variants in the human hypothalamus. **a**, Schematic of cluster-specific scATAC peaks analyses used to prioritize obesity-associated SNPs. **b-c**, Luciferase assays in mouse hypothalamus neuronal cells for sequences that overlap obesity-associated SNPs (**b**) and the reference or obesity-associated SNPs (**c**). pGL4.13 (Promega) with an SV40 early enhancer was used as a positive control and the pGL4.23 empty vector as a negative control. **d**, Genomic snapshot of the *MC4R* locus and two identified enhancer regions with logo plots showing TFBS inference due to the obesity-associated SNP, rs17782313 **e**, Genomic snapshot of the *FTO* locus along with the *IRX3* and *IRX5* genes and the *FTO* enhancer with logo plots showing TFBS inference of HOXD10 due to the obesity-associated SNP, rs8043757. **f**, qRT-PCR of *MC4R* (left) in iPSC and differentiated neurons and control and two *MC4R* enhancer KO cell line (right). Data are represented as mean ± standard deviation, (***) P-value ≤0.001 **g**, qRT-PCR of *FTO, IRX3*, and *IRX5* in human astrocytes treated with CRISPRi and different gRNA targeting the *FTO* enhancer. Data are represented as mean ± standard deviation, (*) P-value≤0.001 (***) P-value≤0.001.

We next tested the enhancer activity of 18 obesity-associated scATAC-seq peaks that overlapped with the strongest obesity-associated SNPs. These peaks are found to be specific to cell clusters, including OPC, MFO, astrocytes. We cloned these sequences into an enhancer assay vector that contains a minimal promoter followed by the luciferase reporter gene and transfected them into mouse neuronal hypothalamus cells. We found 10 out of 18 (55.5%) sequences to have luciferase activity that was significantly higher than the empty vector negative control (**Fig.5b**). These include sequences near the genes *ADCY3, FAIM2, FTO, GIPR, GNAT2, GPRC5B, LEMD2, MC4R, SDCCAG8, SH2B1, SULT1A1, TDH*, and *VPS45*. (**Fig.5b**). Using a similar assay, we also tested the enhancer activity of sequences from scATAC astrocyte-specific peaks (*FTO, SDCCAG8, LPP*) in primary human astrocytes, obtaining similar results to the neuronal cells (**Extended Data Fig.5a**). We next used site-directed mutagenesis to introduce the obesity-associated SNPs to the sequences that showed enhancer activity and tested them for differential activity compared to the unassociated variants. We found that 5 out of the 10 sequences, near the genes *FAIM2, FTO, GIPR, MC4R* and *VPS45* showed significant differential enhancer activity with the obesity-associated variant/s compared to the reference variant (**Fig.5c**).

The SNP that showed differential enhancer activity for *MC4R*, is the second top obesity-associated GWAS SNP and also happens to be the lead SNP, rs17782313. It is located 187k bp from *MC4R* in a sequence that is not conserved in mice. Using TRANSFAC^110^, PROMO^111^, and JASPAR^112^, analysis of the risk allele found it to potentially disrupt binding of GATA binding protein 1 (GATA1), which was shown to bind to the 5’UTR of the *MC4R* gene^113^(**Fig.5d**). To characterize its function and test whether it regulates *MC4R*, we set out to knockout this sequence in neurons using CRISPR-Cas9. As *MC4R* is not expressed in many cell types, we first differentiated WTC11-NGN2 iPSC cells^114^ to functional glutamatergic neurons (see **Methods**) and tested whether it is expressed in these cells. We found *MC4R* to be significantly expressed in these cells (∼8-fold compared to iPSC cells; **Fig.5f**). We then transfected these iPSC cells with two gRNAs targeting both ends of the postulated *MC4R* enhancer along with Cas9 protein. Single-cell colonies were FACS-isolated and screened by genotyping for homozygous cells, finding two colonies. These colonies, along with wild-type cells, were differentiated to neurons and analyzed for *MC4R* expression via qRT-PCR. We found that the two independent *MC4R* enhancer knockout cells lines had significantly lower *MC4R* expression (60%) compared to wild-type cells (**Fig.5f**). Since there are no other genes in the *MC4R* TAD boundary, we did not measure expression for other genes. Taken together, these results showcase that the obesity associated *MC4R* SNP reduces enhancer activity, and that removal of this enhancer leads to reduced *MC4R* expression, suggesting that it regulates this gene.

We next examined the function of the *FTO* enhancer. This sequence was previously shown to regulate its neighboring genes, Iroquois homeobox 3 and 5 (*IRX3*/*5*), in the hypothalamus^115^ and adipocytes^116^. TFBS analysis for the obesity-associated SNP, rs8043757, suggested that it decreases binding affinity for the high mobility group forkhead box P3 (*FOXP3*), which was shown to regulate gonadotropin expression in pituitary in mice^117^(**Fig.5e**). To further characterize its target genes in a cell-type specific manner, we attempted to delete the *FTO* enhancer in astrocytes. However, cells failed to proliferate and form colonies, likely due to the known role of *FTO* in the proliferation of neuronal cells^118^. We thus used CRISPR inactivation (CRISPRi) instead. We designed four gRNAs targeting the identified enhancer and tested them for their ability to downregulate expression by co-transfecting human astrocytes with a nuclease deficient Cas9 (dCas) fused to the KRAB transcriptional repressor. Examination by qRT-PCR found *FTO* to have significantly lower expression for three of the four gRNAs, with the lowest reduction for gRNA-4 (**Fig.5g**). We also analyzed the expression of *FTO* enhancer target genes, *IRX3* and *IRX5*, finding both to have decreased expression levels with all gRNAs, with the highest impact for gRNA-4 (**Fig.5g**). In line with the enhancer being specific to astrocytes, we found that both *IRX3* and *IRX5* were highly expressed in astrocyte populations (**Extended Data Fig.5b**). In summary, we found obesity-associated SNPs located in the enhancer regions of *MC4R* and *FTO* could potentially disrupt binding of TFs to these enhancers, lead to lower enhancer activity and subsequently lower expression of these obesity-associated genes. Combined, this work showcases the ability of integrative scRNA and scATAC-seq to identify gene regulatory elements whose alteration is associated with hypothalamus related phenotypes, such as obesity.

## Discussion

In this study, we utilized integrative single-cell sequencing to characterize mouse and human hypothalamus. Integrative single-cell analysis allowed us to identify various neuronal and non-neuronal cell types, uncover cell-specific regulatory elements and their target genes, and elucidate the transcriptional regulation networks of these cells in the hypothalamus. This atlas identified numerous non-neuronal and neuronal cells that play key roles in controlling hypothalamus-associated physiological processes. By using both female and male hypothalamus, we were able to identify many sex-differential expressed genes and regulatory elements. This integrative data also allowed us to identify and validate cell-type specific regulatory elements that encompass obesity-associated SNPs, including *FTO* and *MC4R*, which alter their enhancer activity. Deletion of the obesity associated *MC4R* enhancer or CRISPRi targeting of the *FTO* enhancer in astrocytes validated their target genes.

Using gene markers from previous scRNA-seq studies^29-41^, we identified many neuronal cell populations in mice (N=33), including glutamatergic, GABAergic, specialized neuron clusters, OPC, ODC, astrocytes, and microglia, with the majority of cells (90%) being neurons. We were also able to identify four specialized neuron cell types that were not identified in these studies. In contrast, in the human hypothalamus, neurons made up less than 5% of the total cells while the majority of cells were oligodendrocyte associated. This is in line with two recent studies in which multiomic single-cell sequencing was performed using post-mortem human midbrain or frontal lobe, showed similar recovery of neuronal and oligodendrocyte populations^63,64^. Similarly, the recent study by Siletti et al,. in which ∼3.4 million nuclei from 106 sections of three human adult brains were sequenced, observed a large number of oligodendrocytes (∼600K) despite the fact that they isolated neurons by FACS with NeuN (similar to what was done here) aiming to collect 90% neurons and 10% non-neuronal cells^119^. This might be due to human neurons being sensitive to nuclear permeabilization or samples being collected post-mortem when nucleus neurons are the most sensitive to hypoxia^120^ as well as the age of donors (**Supplemental Table 1**).

Tissue- or cell-specific gene regulatory networks control cell fate specification and drive dynamic processes, such as cell differentiation. Integrative single-cell sequencing allowed us to identify regulatory elements and link them to their regulated genes in specific clusters and further map TFs activity in a cell type-specific manner. For example, in the mouse hypothalamus, we identified two regulatory elements linked to *Pomc* expression only in the *Pomc*/*Cartpt* neurons which have been reported to be enhancers of this gene^59^. In the human hypothalamus, we identified four regulatory elements linked to *GAD1* expression, one of which is located ∼50kb from its promoter and has already been characterized as its enhancer^121^. We also identified numerous cell-specific scATAC peaks linked to expressions of cell marker genes, including *SLC32A1* in GABAergic neurons, *SLC16A7* in glutamatergic neurons, *TH* and *DDC* in dopamin neurons, and *PDGFRa* in OPC. These regulatory elements are all novel and can provide deep understanding of the transcriptional regulation of these cell type-specific marker genes. Furthermore, by performing TFBS enrichment analyses on scATAC peaks, we identified many cell type-specific networks of TF activity. Comparison of cell-types across mice and human hypothalamus allowed us to identify shared core TFs that could serve as cell cluster-specific regulatory machinery. We also found genes that are regulated by these core TFs and had cell type-specific functions, further confirming that each cell type requires distinctive gene regulatory networks to regulate its function and activities.

We identified numerous genes that are sex-differentially expressed across various cell clusters in mouse and human hypothalamus. We found that sex effects are small, but ubiquitous across cell populations. When examining neuropeptides, we found several neuropeptide genes that were sex-differentially expressed, some of which were previously reported. For example, we found *NPY* to be more highly expressed in male glutamatergic neurons compared to females, similar to studies showing *NPY* brain expression levels are lower in females than males, contributing to higher anxiety and anxiogenic phenotype in males^122,123^. Interestingly, we also found novel sex-differentially expressed neuropeptides, such as *HCRT* in glutamatergic neurons, T*RH* in GABAergic neurons. *HCRT* was shown to have sex-differential effects on body weight and body composition when ablated in mice, however, its expression was not reported^124^. This sex-differential effects might be due to its sex-differential expression levels in the hypothalamus. Similarly, *TRH*, which stimulates the release of thyroid stimulating hormone (*TSH*) and prolactin from the anterior pituitary^67^, was not shown to be differentially expressed between sexes in GABAergic neurons. Furthermore, we found sex-specific regulatory elements that are linked to sex-differentially expressed genes. For examples, in GABAergic neurons, we found a male-specific scATAC peak was linked to the expression of proenkephalin (*PENK*), which was shown to be involved in pain perception and response to stress in males rats^105^. Sex-dimorphic expression of some of these key neuropeptides might be involved in sex-differential processes, such as reproductive and social behaviors, and physiological responses to environment cues.

Our integrative single-cell datasets allowed the identification of novel genes, regulatory elements and pathways associated with hypothalamus related phenotypes. Here, as an example, we showcase the advantage of this approach to identify obesity-associated regulatory elements. Numerous obesity GWAS loci reside near hypothalamus expressed genes^9,10^, several of which are associated with the hypothalamic leptin-melanocortin system^10^, and have been difficult to detect due to the inability to identify gene regulatory elements in distinct neuronal subpopulations. We were able to identify multiple hypothalamus cell type specific scATAC peaks that overlap with obesity-associated SNPs, including the top two obesity associated loci, *FTO* and *MC4R*. This allowed us to pinpoint the cell types in which these potential obesity-associated regulatory elements are involved, finding them to be distributed in various cell populations, including OPC, ODC, astrocyte, and neurons. Interestingly, we found no potential obesity-associated enhancers in the microglia population. Utilizing enhancer assays, we found that more than half of these overlapping scATAC peaks have enhancer activity in either neurons or astrocytes. Additional cell types and or primary cells could also be used to dissect the function of these sequences. Amongst the active sequences, we found that half of them altered enhancer activity. Interestingly, in all these cases this led to a reduction rather than an increase in enhancer activity. Our CRISPR experiments for *MC4R* and *FTO* in iPSC-derived neurons and primary astrocytes further confirmed the target genes of these enhancers, showcasing the utility of these datasets.

## Methods

### Nuclei isolation and library preparation

For mice, the whole hypothalamus of three male and three female 6 weeks-old *Mc4r*^*t2aCre/t2aCre*^ *x Gt(ROSA)26Sor*^*tm2(CAG-NuTRAP)Evdr*^ C57BL/6J mice generated by Dr. Vaisse^44^ were subjected to nuclei isolation following the 10X Genomics established protocol (CG000375-Rev A). In brief, flash-frozen hypothalamus samples were lysed and homogenized in 0.1% NP40 lysis buffer supplemented with RNase inhibitor (Sigma, 3335402001). The nuclei mixture was filtered through a 70um strainer and then stained with 7-AAD (7-aminoactinomycin D) in PBS with 1% BSA and RNase inhibitor. Nuclei were FACS-sorted using the ARIA Fusion Cell Sorter with a 100um nozzle. Nuclei were washed and pre-metallized with 0.01% digitonin lysis buffer supplemented with RNase inhibitor. For each sample, 20,000 nuclei were subjected for GEX and ATAC libraries prepared using Chromium Next GEM Single Cell Multiome ATAC + Gene Expression (CG000338 Rev B). Both libraries were sequenced using HiSeq 4000 with 25,000 paired reads per nucleus. For human samples, three male and three female hypothalami were collected post-mortem at UCSF medical center following the UCSF human research protection program institutional review board protocol number 21-34261. The donors’ age ranged from 43-82 (**Supplementary Table 1**). The human samples were subjected to the same protocols as mouse hypothalamus. Due to the large size of the human sample, the hypothalamus was split in half and subjected to the lysis step and pooled together prior to FACS. 40,000 nuclei per sample were used for the library prep.

### Data preprocessing and quality control

Demultiplexed scRNA- and scATAC-seq fastq files were generated using CellRanger Arc mkfastq (10x Genomics, 7.0.1). scRNA and scATAC-seq data were aligned to pre-built reference genomes GRCh38 and GRCm38 from 10x Genomics. Barcoded count matrices of gene expression and ATAC data were generated using CellRanger ARC pipeline (version 2.0.0) (10X Genomics). Count and peak matrices and fragment files from CellRanger were analyzed using the ‘Seurat’ package (version 4.1.1)^125^. Quality cells were selected using the following quality control metrics: TSS enrichment greater than 1; nucleosome signal greater than 2; fraction of mitochondrial genes less than 30%; between 100 and 7500 genes detected in each cell; and between 1000 and 30000 peaks with at least one readcount detected in each cell. For the human dataset, contaminating ambient RNA was detected using SoupX and the corrected count matrix used for downstream analysis. Gene annotations from Ensembl Database v79 (mouse) and Ensembl Database v86 were used.

### scRNA-seq data analysis

scRNA-seq data of nuclei were analyzed using ‘Seurat’ after quality control filtering. Gene expression data were log-normalized and multiplied by 10,000 using the NormalizeData and ScaleData functions. The top 3,000 variable features were identified using FindVariableFeatures function. Batch-correction between samples was performed using Harmony^126^. Dimensionality reduction was performed with PCA and the top 50 principal components were kept. Nearest neighbors used Harmony embeddings, and clustering was performed using resolution = 0.6. Differentially expressed genes were identified using^127^ Seurat’s FindAllMarkers function^125^, using Wilcoxon Rank Sum test, filtering for a minimum log2 fold change threshold of 0.25, and minimum fraction of 0.1 cells in the population.

### scATAC-seq data analysis

scATAC-seq data were processed using ‘Signac’ R package (version 1.7.0). We apply term frequency-inverse document frequency normalization to ATAC peaks, followed by feature selection and dimensionality reduction using singular value decomposition. Batch-correction was performed using Harmony. 50 dimensions for ATAC and 50 dimensions RNA PCA dimensions were used to construct the weighted nearest neighbor graph. Peak calling was performed using MACS2^128^, and peaks not found on standard chromosomes and within blacklist regions were pruned from the peak set. Differentially accessible regions were identified using FindAllMarkers function, using likelihood ratio test, filtering for a minimum log2 fold change of 0.25, minimum fraction of 0.05 cells in the population, including number of ATAC counts as latent variable. Linked peaks were identified using Signac’s LinkPeaks function using peaks found in at least 3 cells, distance cutoff of 1 million basepairs, p-value cutoff of 0.05. Nearest genes to scATAC-seq peaks were determined using ClosestFeatures function.

### Motif enrichment analysis

Motif analysis was performed on accessible peaks using ChromVar^72^. Motif position weight matrices were downloaded from the JASPAR 2020 database. Differential testing on chromVar Z-scores was performed using the FindAllMarkers function (Wilcoxon Rank Sum test) in Signac with minimum fraction of cells as 0.1. Peaks containing motifs were identified using the ‘motifmatchr’ version 1.18.0^129^. GO terms were identified using GENEontology.

### SNPs selection and overlapped with cell type-specific ATAC peaks

Indexed SNPs associated with obesity were collected from the NHGRI-EBI GWAS Catalog^107^. LD expansion was then performed using LDLink^130^ to identify LD linked SNPs with r2>=0.8. LD linked SNPs were filtered to exclude variants in coding regions based on annotations in dbSNP to obtain a final set of noncoding SNPs. The coordinates of these SNPs were intersected with differentially accessible ATAC peaks. P-values for SNP enrichment in ATAC peaks were computed using the binomial test as implemented in SciPy. We set the number of successes as SNPs that overlapped the ATAC peaks. We set the number of trials as number of SNPs from obesity collected as described above. The coordinates of these SNPs were intersected with differentially accessible ATAC peaks, found using Seurat’s ‘FindAllMarkers’ function using likelihood ratio test with log2fold change threshold > 0.15. ‘motifbreakR’ package^131^ was used to predict effects of SNPs on motifs on its surrounding region.

### Luciferase assay

The scATAC-seq peaks that overlapped obesity-associated SNPs were PCR amplified from human genomic DNA (see primers in **Supplementary Table 2**) and cloned into the pGL4.23 plasmid (Promega, E84111). The associated SNPs were then introduced into these plasmids by PCR amplification with primers containing the associated variants. The PCR products were then treated with DpnI to remove WT and the constructs with the associated SNPs were confirmed with Sanger sequencing. The These unassociated and associated constructs (200ng) along with Renilla luciferase (20ng), to correct for the transfection efficiency, were transfected into mouse hypothalamus Pomc neuronal cells (mHypoA-POMC/GFP-1 from Sanbio #CLU500) or human astrocytes (CCF-STTG, ATCC) grown in 96-well plates of ^132^ using X-tremeGene (Sigma, 6366244001) following the manufacturer’s protocol. An empty pGL4.23 vector was used as negative control and pGL4.13 (Promega, E668A), which has an SV40 early enhancer, as a positive control. Forty-eight hours post transfection, cells were lysed, and luciferase activity was measured using the Dual-Luciferase Reporter Assay System (Promega, E1910). Six technical replicates were performed for each condition.

### Neuronal differentiation

WTC11 iPSC cells (a generous gift from Li Gan (Gladstone Institute)) were maintained in Matrigel-coated plates with mTeSR1 basal media (StemCell, 85850). Cells were subcultured using Accutase and plated with mTeSR1 media supplemented with 10uM Rock inhibitor (Fisher Scientific, NC1286855). To differentiate these cells to mature neurons, we followed the previously published protocol in ^114^. In brief, cells were plated in Matrigel-coated 6-well plate in pre-differentiation media (see recipe in **Supplementary Table 3**) and Rock inhibitor. Media was changed every day for the next 2 days. On day 4, cells were then sub-cultured into poly-L-Ornithine-coated plates (Sigma, P3665) in maturation media (**Supplementary Table 3**) supplemented with 2ug/mL doxycycline (Sigma, D3447). Half of the media was then replaced with maturation media after 7 days. Cells were collected at day 14 of differentiation.

*MC4R* enhancer knockout cells were generated by transfecting WTC11-NGN2 iPSC cells in 6-well plates with 6.25ug Cas9 protein (Fisher Scientific, A36498) and 800ng sgRNAs (IDT), and 0.5ug GFP plasmid (Addgene, 13031) using Lipofectamine CRISPRMax Cas9 transfection reagent (Fisher Scientific, CMAX00-003) following the manufacturer’s protocol. After 48 hours, GFP+ cells were isolated into 96 well-plates into single clones using FACS (BD FACSAria Fusion) (see gRNAs in **Supplementary Table 2**). These colonies were then genotyped by PCR with primers flanking 200bp the deletion site. The knockout genotype has 400 bp PCR product while WT has 3500 bp PCR product (see primers in **Supplementary Table 2**). Cells were then subjected to the neuron differentiation protocol described above.

### RNA Isolation, cDNA synthesis and qRT-PCR

Total RNA was extracted from cells using RNeasy Plus kit (Qiagen, 74106). Reverse transcription was performed with 1μg of total RNA using qScript cDNA Synthesis Kit (Quantabio, 95047) following the manufacturer’s protocol. qRT-PCR was performed on QuantStudio 6 Real Time PCR system (ThermoFisher) using Sso Fast (Biorad, 1705205). Statistical analysis was done using ddct method with GAPDH primers as control (see primer sequences in **Supplementary Table 2**). Gene expression results were generated using mean values for over 4-6 biological replicates.

### CRISPRi study

gRNAs targeting the *FTO* enhancer were designed using CRISPick (Broad Institute)^133^ (see gRNAs in **Supplementary Table 2**). gRNAs were cloned into an AAV vector (pAAV-U6-sasgRNA-CMV-mCherry-WPREpA) and co-transfected into human astrocyte cells (CCF-STTG1, ATCC) along with dCas9-KRAB (kind gift from Dr. Alejandro Lomniczi OHSU). After two days, cells were lysed, and RNA and cDNA were prepared as mentioned above.

## Acknowledgments

We would like to thank Dr. Alejandro Lomniczi for kindly providing the pCVM-sadCas9-KRAB vector. This work was funded in part by the National Institute of Diabetes and Digestive and Kidney Disease (NIDDK) R01DK116738 (C.V. and N.A), R01DK106404 and R01DK060540 (CV), the UCSF Nutrition and Obesity Research Center P30 DK098722, the California Institute for Regenerative Medicine (CIRM) postdoctoral fellowship (H.P.N.), the UCSF Hillblom Center for the Biology of Aging and Bakar Aging Research Institute Graduate Fellowship (C.S.Y.C), a Societe Francaise du Diabete and a fondation Servier fellowship (FM), Vera Long Fellowship (AB), and start-up funds from the Evergrande Center (M.H).

## Author contributions

H.P.N., N.A. and CV conceived and designed the study. H.P.N., D.L.R., R.S., L.H., M.N., A.U., C.B., K.A. performed experiments., H.C., C.S.Y.C., M.G.G., M.H. analyzed genomic data, F.M., A.D., and C.V. provided mice hypothalami, E.H. provide human samples, H.P.N., C.S.Y.C. and N.A. wrote the manuscript with contributions from all authors.

## Competing interests

N.A. is a cofounder and on the scientific advisory board of Regel Therapeutics. N.A. receives funding from BioMarin Pharmaceutical Incorporate.

## Data and code availability

Raw and processed data from this study have been deposited in the NCBI short read archive (SRA) as SRA12255727and SRA12255698. Code used for scRNA-and scATAC-seq analysis is available at https://github.com/candacechan/sc_hypothalamus.

---

**Extended Data Fig.1.**
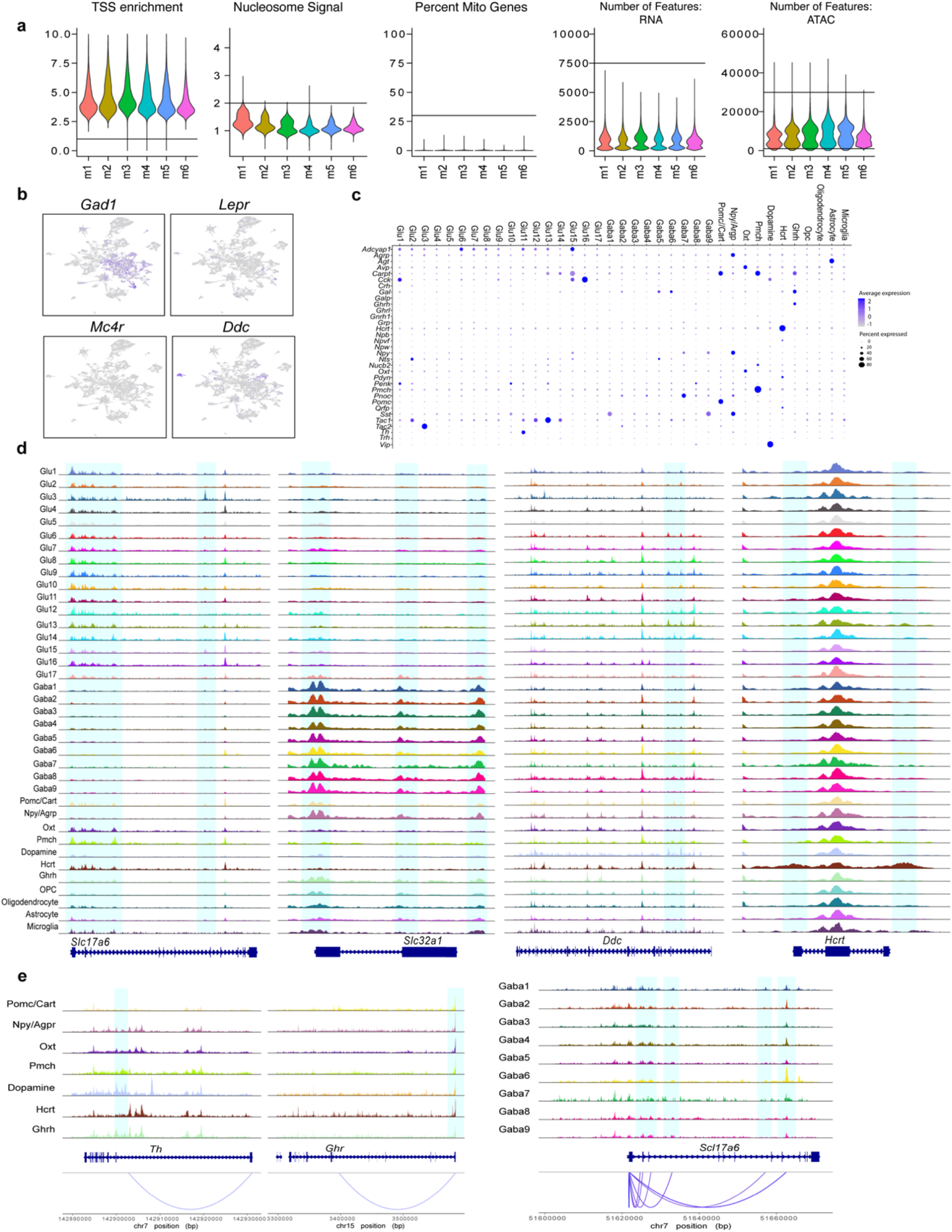
Combined scRNA and ATAC profiling of the mouse hypothalamus. **a**, Violin plot of quality control metrics: TSS enrichment scores, nucleosome signal, percentage of mitochondrial genes, number of features in RNA data, and number of features in ATAC data. **b**, Uniform Manifold Approximation and Projection (UMAP) of scRNA of *Gad1, Lepr, Mc4r*, and *Ddc*. **c**, Dot plot of neuropeptide gene expression across all cell types in mouse hypothalamus. **d**, Coverage plot of genes, *Slc17a6, Slc32a1, Ddc, Hcrt* showing scATAC peaks highlighted in blue around the gene body. **f**, scATAC peaks to gene linkage plots for *Th* and *Ghr* in specialized neurons, and *Slc17a6* in the GABAergic neurons with the genome tracks. The linked scATAC peaks to gene promoters are highlighted in blue.

**Extended Data Fig.2.**
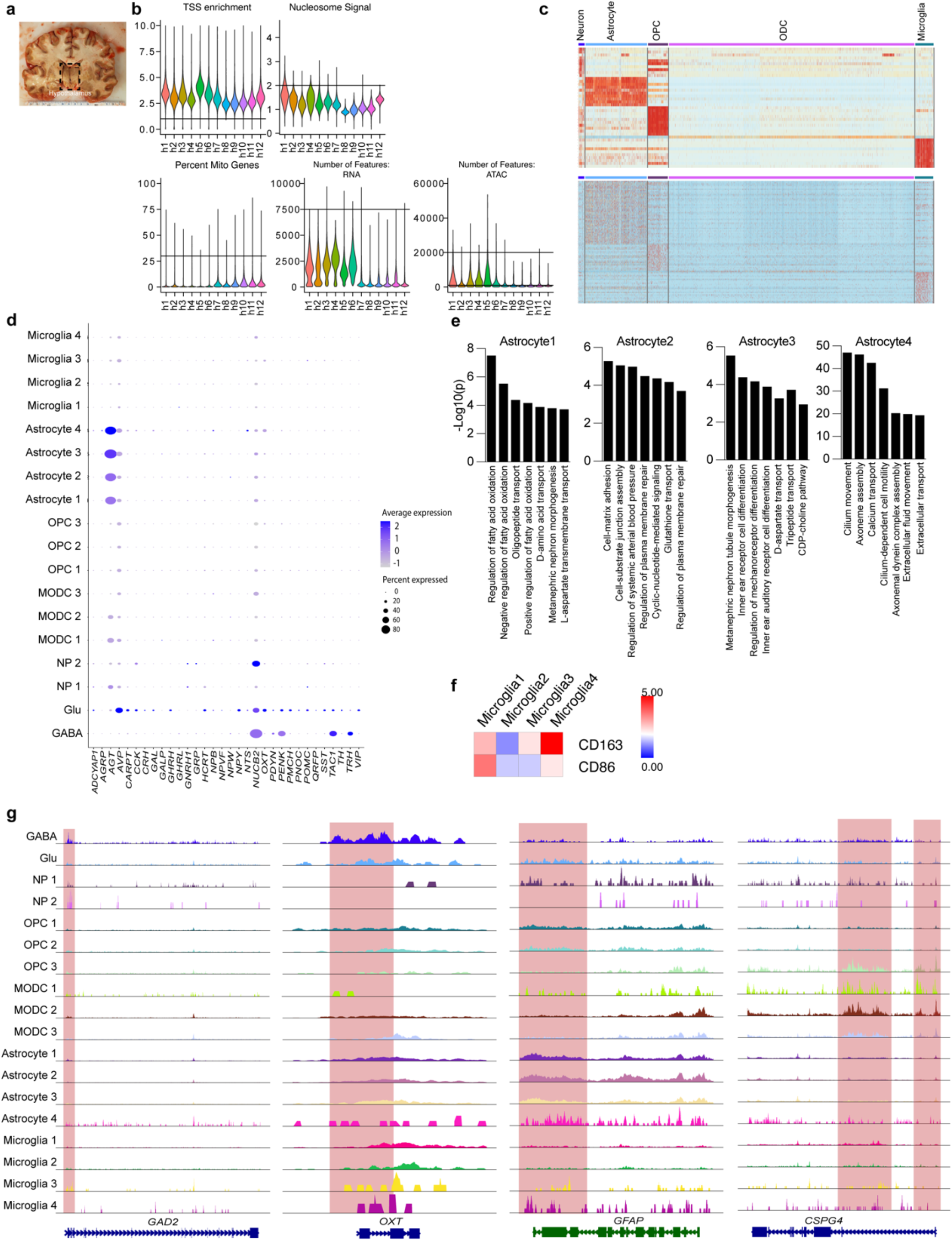
Combined scRNA and ATAC profiling of the human hypothalamus. **a**, Representative image of human hypothalamus section. **b**, Violin plot of quality control metrics: TSS enrichment scores, nucleosome signal, percentage of mitochondrial genes, number of features in RNA data, and number of features in ATAC data. **c**, Heatmap of the top five differentially expressed genes and the top ten scATAC peaks across all cell populations in human hypothalamus prior to subclustering. **d**, Dot plot of neuropeptide gene expression across all cell types in the human hypothalamus. **e**, Gene ontology analyses of all astrocyte cell clusters. **f**, Heatmap of *CD163* and *CD86* expression in astrocyte populations. **g**, Coverage plot of genes, *GAD2, OXT, GFAP, CSPG4* showing scATAC peaks highlighted in blue around the gene body.

**Extended Data Fig.3.**
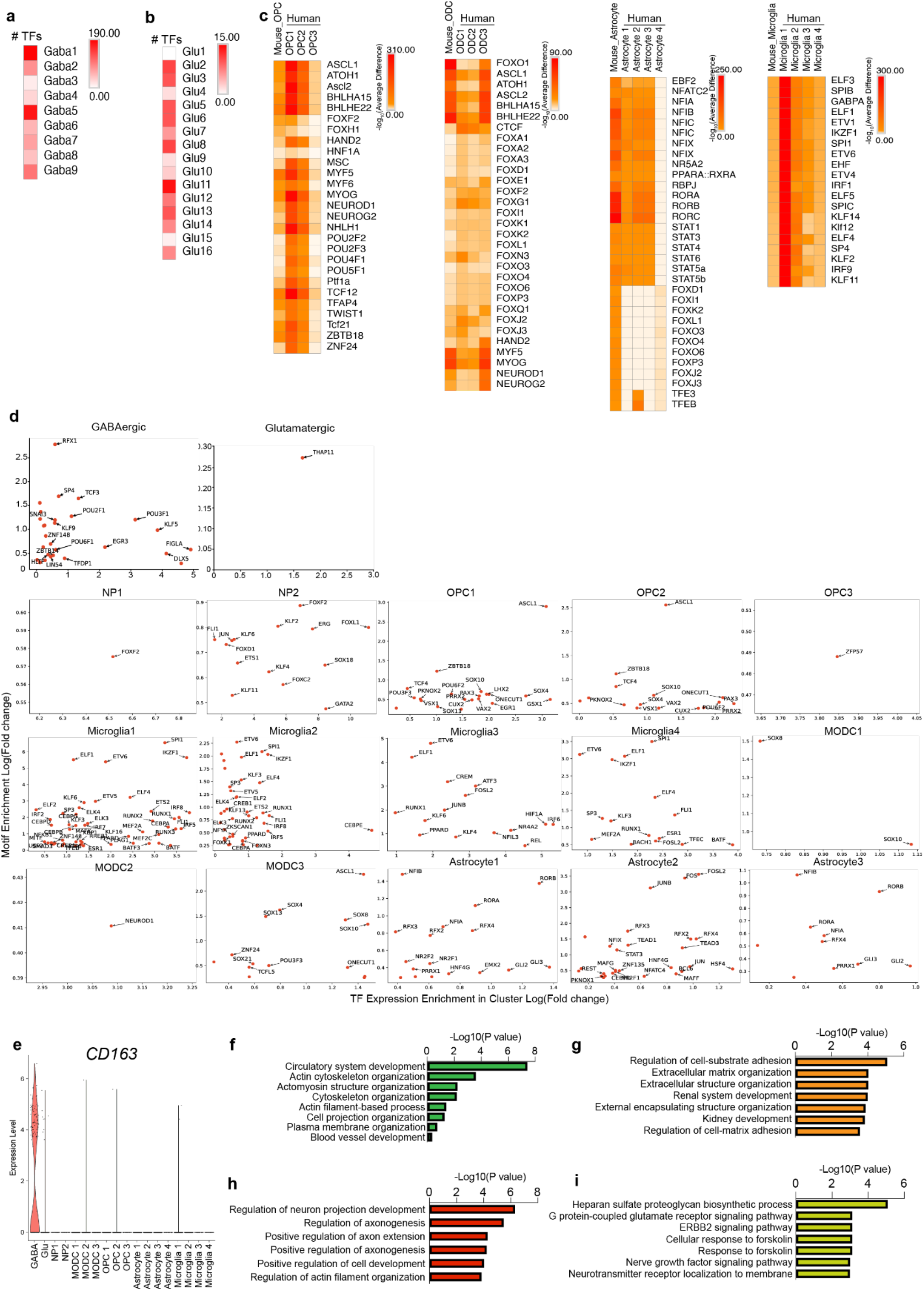
TFs and TFBS enrichment in mice and human hypothalamus cell populations. **a**, Heatmap showing the number of TFs in mouse GABAergic neurons shared with human GABAergic neurons. **b**, Heatmap showing the number of TFs in mouse glutamatergic neurons shared with human GABAergic neurons. **c**, Heatmap of TFBS in OPC, ODC, astrocyte, and microglia populations shared by mice and human. **d**, TFs that show both upregulation of gene expression and motif enrichment in all populations. **e**, Violin plot of *CD163* expression. **f-i**, Gene ontology of genes regulated by mouse-specific TFs in GABAergic (**f**), glutamatergic (**g**) neurons, and human-specific TFs in GABAergic (**h**) and glutamatergic (**i**) neurons.

**Extended Data Fig.4.**
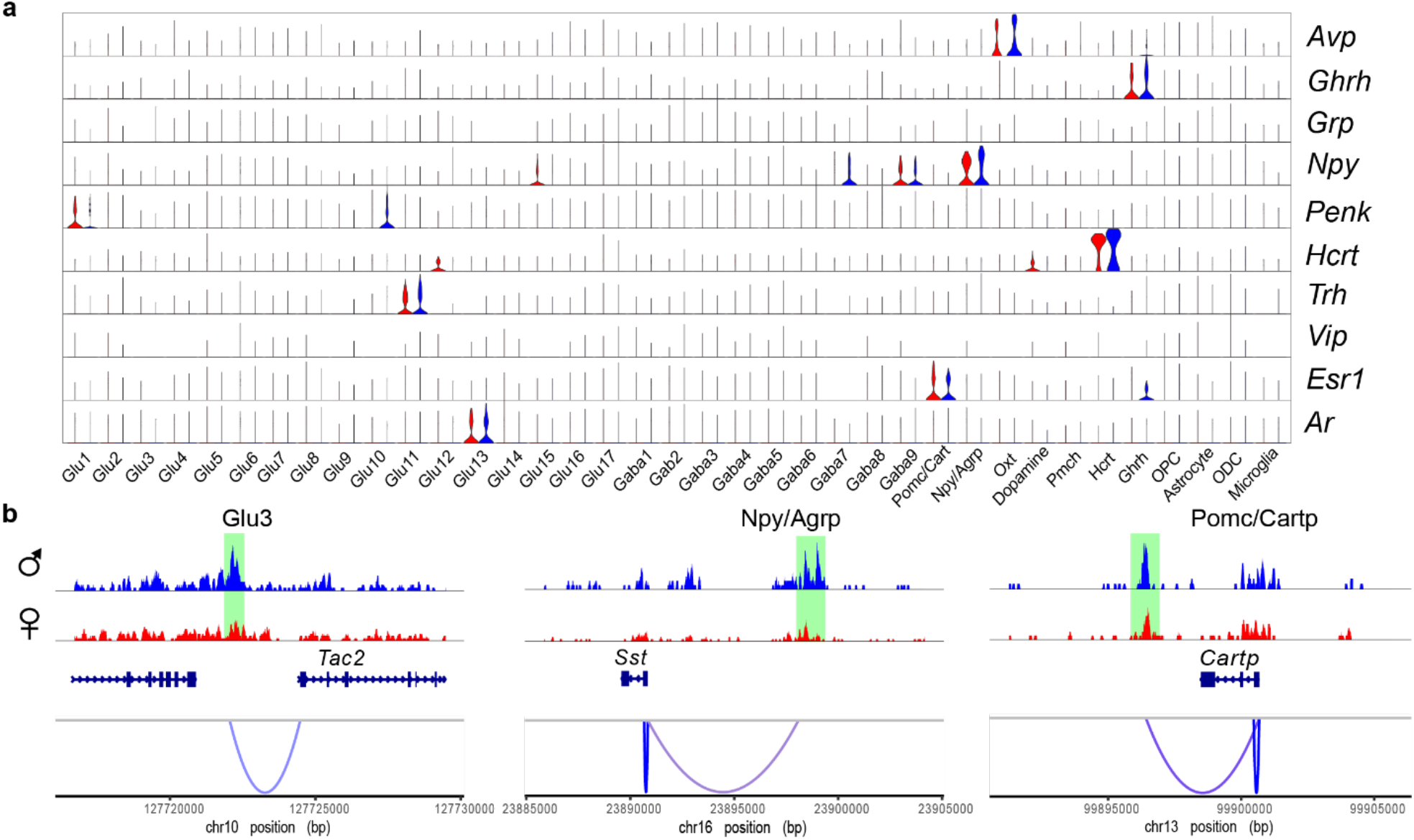
Sex-differential expression and regulation. **a**, Violin plots of sex-differentially expressed neuropeptide genes in mouse hypothalamus. **b**, Linkage plots showing sex-specific scATAC peaks linked to sex-differentially expressed genes, including *Tac2* in Glu3 cluster, *Sst* in Npy/Agrp neurons, and *Cartpt* in Pomc/Cartp neurons.

**Extended Data Fig.5.**
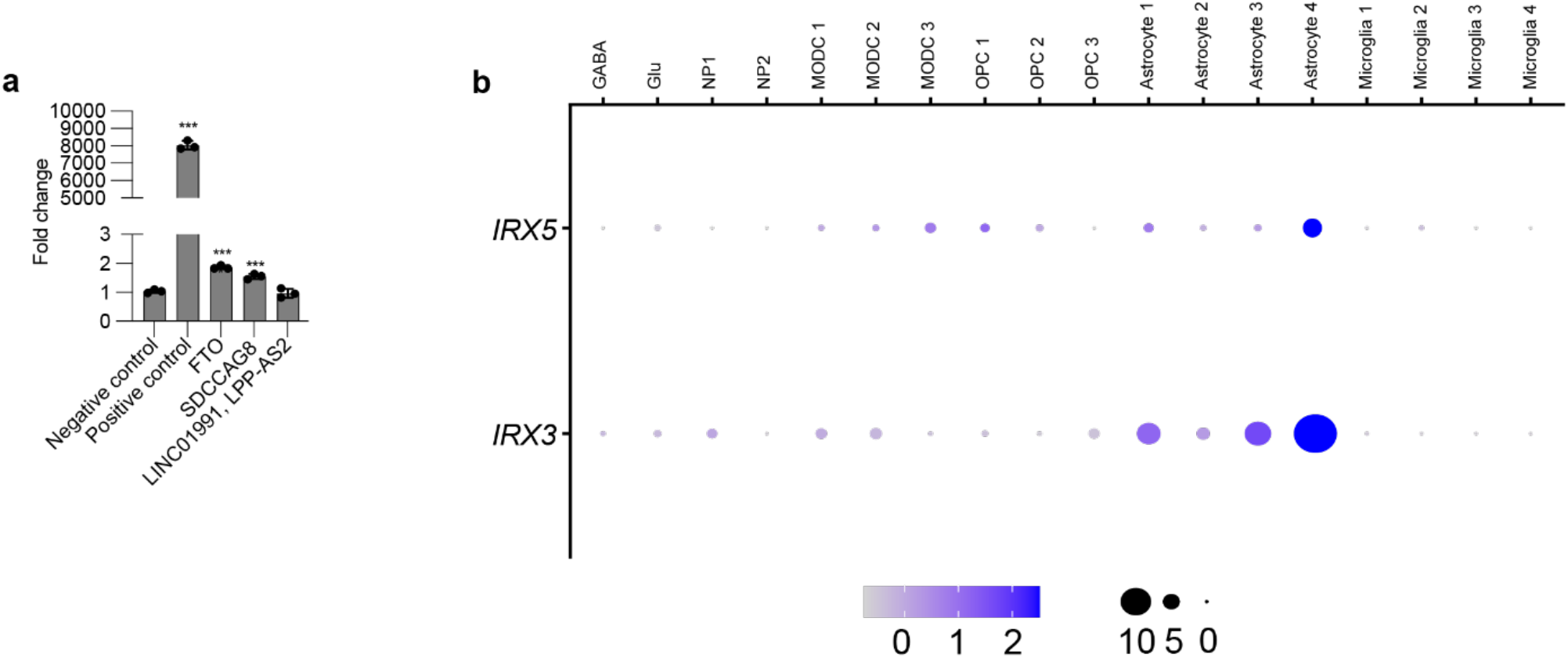
**a**, Luciferase assays in human astrocytes for astrocyte-specific scATAC-seq peaks overlapping obesity-associated SNPs. **b**, Dot plots of *IRX3* and *IRX5* expression across cell populations in the human hypothalamus.

**Supplementary Table 1.** Human hypothalamus sample information.

| **ID** | **Age/Gender/PMI** | **Regions** |
| --- | --- | --- |
| **Hypo 1** | 82/male/7hr | Hypothalamus (between optic chiasm and mammillary body) |
| **Hypo 2** | 62/male/7hr | Hypothalamus (between optic chiasm and mammillary body) |
| **Hypo 3** | 62/male/10hr | Hypothalamus (between optic chiasm and mammillary body) |
| **Hypo 4** | 43/female/12hr | Hypothalamus (between optic chiasm and mammillary body) |
| **Hypo 5** | 45/female/4hr | Hypothalamus (between optic chiasm and mammillary body) |
| **Hypo 6** | 73/female/8hr | Hypothalamus (between optic chiasm and mammillary body) |

**Supplementary Table 2.**
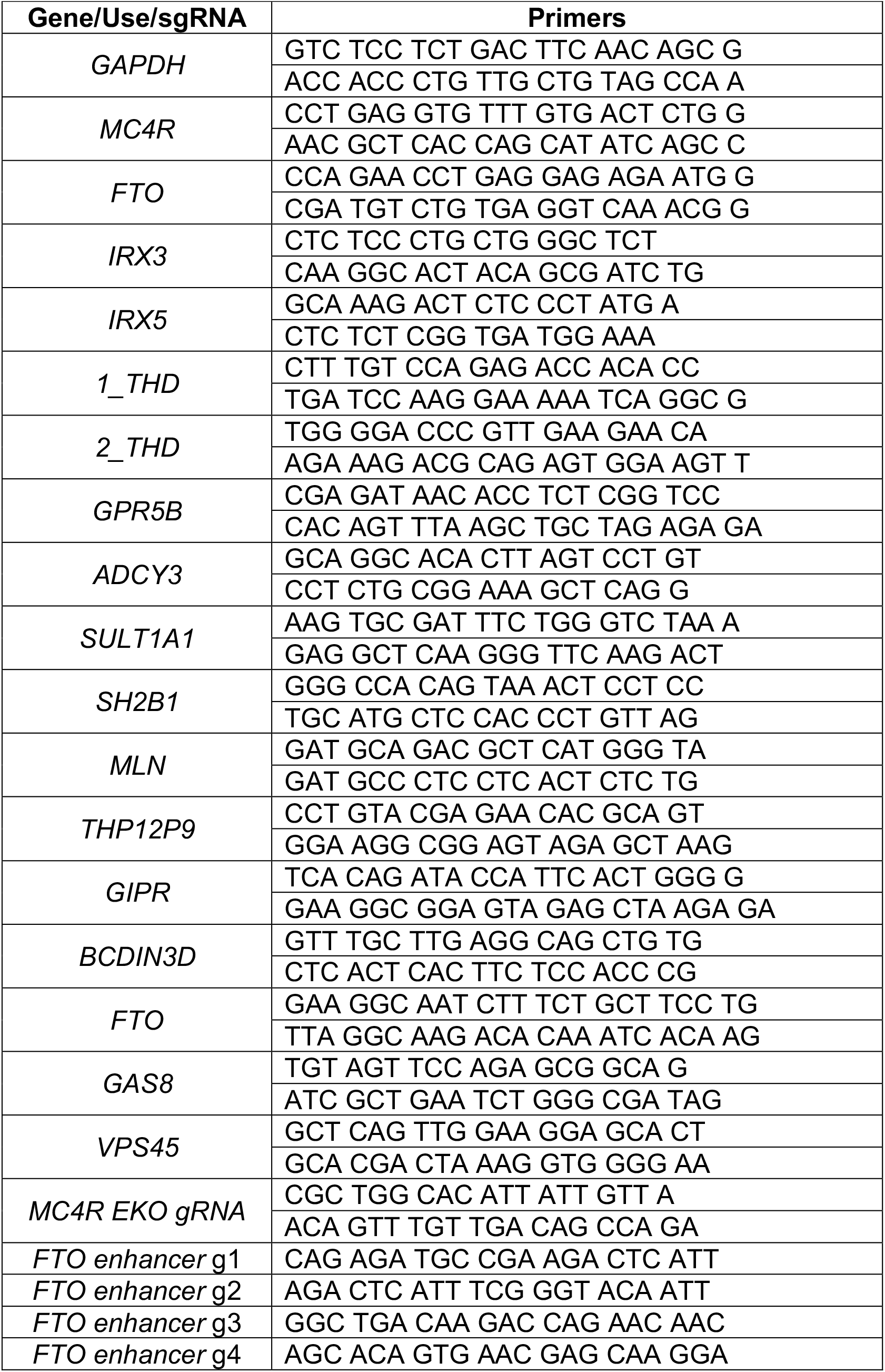
Sequences of primers for RT-qPCR, cloning, genotyping and gRNAs.

**Supplementary Table 3.** Predifferntiation media and maturation media for neuron differentiation from WTc11 iPSC cells.

| Predifferentiation media |  |  |  |  |
| --- | --- | --- | --- | --- |
| Reagent | Stock | Final conc. | Example (50mL) | Example (10mL) |
| KnockOUT DMEM/F-12 | -- | 100% | 49mL | 10mL Pre-differentiation media |
| N-2 Supplement (100X) | 100X | 1X | 0.5mL |  |
| NEAA (100X) | 100X | 1X | 0.5mL |  |
| BDNF | 25 ug/mL | 10 ng/mL | **Add these right before use** | 4uL |
| NT-3 | 25 ug/mL | 10 ng/mL |  | 4uL |
| Laminin | 1.2 mg/mL | 1.2 ug/mL |  | 10uL |
| Doxycycline | 1 mg/mL | 2 ug/mL |  | 20uL |
| Maturation media |  |  |  |  |
| Reagent | Stock | Final conc. | Example (50mL) | Example (10mL) |
| Neuralbasal-A Medium | -- | 50% | 24.25mL | 10 mL Maturation Medium |
| DMEM/F-12, HEPES | -- | 50% | 24.25mL |  |
| NEAA(100x) | 100X | 1X | 0.5mL |  |
| B-27 Supplement (50x) | 50X | 0.5X | 0.5mL |  |
| N-2 Supplement (100x) | 100X | 0.5X | 0.25mL |  |
| GlutaMAX Supplement (100x) | 100X | 0.5X | 0.25mL |  |
| BDNF | 25 ug/mL | 10 ng/mL | **Add these right before use** | 4uL |
| NT-3 | 25 ug/mL | 10 ng/mL |  | 4uL |
| Laminin | 1.2 mg/mL | 1.2 ug/mL |  | 10uL |

## Notes

### Competing Interest Statement

N.A. is a cofounder and on the scientific advisory board of Regel Therapeutics and receives funding from BioMarin Pharmaceutical Incorporate.

### Summary of Updates

We updated the competing interests statement.

